# The ornithine-arginine cycle supported a toxic, metalimnic *Planktothrix rubescens* bloom

**DOI:** 10.1101/2025.08.04.668501

**Authors:** Brittany N. Zepernick, David J. Niknejad, Emily E. Chase, Blessing A. Abiodun, Meaghan J. Adler, Katelyn A. Houghton, Jason L. Olavesen, Qudus Sarumi, Alexander R. Truchon, Jillian L. Walton, Jack H. Cheshire, Keara Stanislawczyk, Lauren N. Hart, Hans W. Paerl, Justin D. Chaffin, Gregory L. Boyer, Hector F. Castro, Shawn R. Campagna, George S. Bullerjahn, Steven W. Wilhelm

## Abstract

*Planktothrix rubescens* is distinct from other cyanobacterial harmful algal bloom (cHAB) genera: the crimson-red cHAB thrives in the cold, low-light, nutrient-limited metalimnion. Studies have attributed this ecological success to buoyancy regulation, low-light adaptations, and the uptake of nitrogen-rich amino acids. Yet, it remains to be mechanistically determined how this cHAB maintains physiological nutrient quotas in the metalimnion due to limited *in situ* molecular studies. We employed metagenomics and metabolomics to investigate a toxigenic *P. rubescens* bloom in Meads Quarry (Knoxville, TN, USA) observed in two separate years. Our results suggest a perennial, metalimnic *P. rubescens* population may exist, with spring turnover facilitating seasonal migration to the epilimnion. Although *P. rubescens* dominated the epilimnion and metalimnion, intracellular metabolite pools grouped by depth and suggested depth-discrete partitioning of the arginine deiminase-mediated ornithine-arginine cycle (OAC, *i.e.,* urea cycle). While the arginine influx driving the OAC is unclear, we hypothesize this input is provided *via* the uptake of urea or nitrogen-rich amino acids. Further, we demonstrate arginine deiminase (*argE*) is broadly distributed in *Planktothrix* genera and known microcystin producers, suggesting *argE*-mediated arginine recycling *via* the OAC may influence the fitness of toxigenic cHAB genera which require ample nitrogen to synthesize microcystins. Cumulatively, our results serve as a case study to provide insight on the metabolic pathways driving the ecological success of metalimnic *P. rubescens* blooms. On a broader scale, this work strengthens the case that alternative nitrogen metabolism – including urea utilization, amino acid uptake, and the OAC – is a driver of toxigenic cyanobacterial blooms in fresh waters.

## Introduction

Cyanobacterial harmful algal bloom (cHAB) genera such as *Microcystis, Dolichospermum*, and *Aphanizomenon* spp. are well-studied due to their emerald-green surface scums which can dominate warm, high-light, nutrient-replete epilimnions (Paerl and Huisman, 2008; Steffen et al., 2017; Wilhelm et al., 2020). In contrast, the crimson-red cHAB *Planktothrix rubescens* has garnered attention for its success in the cold, low-light, nutrient-limited metalimnions (Babanazarova et al., 2013; Lenard, 2009; Walsby et al., 2006). Indeed, *P. rubescens* has been termed “one of the most studied filamentous cyanoprokaryotes” (Dokulil and Teubner, 2012) due in part to its unusual niche and peculiar ecophysiology.

By various accounts, *P. rubescens* could be classified as atypical amongst cHABs. While many cHAB genera bloom in eutrophic, nutrient-replete freshwaters (Olokotum et al., 2020; Paerl et al., 2011; Steffen et al., 2017), *P. rubescens* is commonly reported in oligotrophic, nutrient-limited fresh waters (Ernst et al., 2009; Jacquet et al., 2005; Suarez et al., 2023). Peak growth and density for most cHAB genera coincide with warm, summer temperatures (> 23° C) (Huisman et al., 2018; Paerl and Huisman, 2008; Visser et al., 2016), yet *P. rubescens* attains peak growth and density at cold, winter-spring temperatures (< 15° C) (Akçaalan et al., 2014; Naselli-Flores et al., 2007; Wagner and Bullerjahn, 2024). While most cHAB genera sink out of the water column to overwinter in sediments as akinetes or vegetative cells (Cirés et al., 2013; Fallon and Brock, 1981; Takamura et al., 1984), *P. rubescens* actively grows and accumulates in the metalimnion (Bright and Walsby, 2000; Micheletti et al., 1998; Walsby and Schanz, 2002), often persisting at high densities (> 10 x 10^6^ cells L^-1^) year-round (Anneville et al., 2015; Naselli-Flores et al., 2007; Walsby et al., 2004). Consequently, various studies have attempted to deduce how *P. rubescens* perseveres in this niche, attributing its success to superior buoyancy regulation (Walsby et al., 1983; Walsby et al., 2004; Walsby et al., 2006), low-light adaptations (Bright and Walsby, 2000; Oberhaus et al., 2007; Selmeczy et al., 2025), and the light-dependent uptake of nitrogen-rich amino acids (Walsby and Jüttner, 2006; Zotina et al., 2003). Yet, it remains to be fully elucidated how this bloom-former maintains physiological quotas long-term in the metalimnion – with *in situ* molecular and metabolomic studies notably lacking.

We performed a molecular investigation of a *P. rubescens* bloom in Meads Quarry, Knoxville, TN (USA). The field experiment was implemented by graduate students at the *University of Tennessee* enrolled in *MICRO 669: Advanced Techniques in Field Microbiology*. Students integrated metagenomic and metabolomic data to distinguish the bloom’s genomic potential and metabolic profile. Depth-discrete analyses revealed a toxic *P. rubescens* bloom spanning the cold (<15 °C), light-replete (∼40 μmol photons m^-1^ s^-2^), nitrate-replete (∼12 μM) epilimnion and the cold (<15°C), light-limited (<1 μmol photons m^-1^ s^-2^), nitrate-replete (∼20 μM) metalimnion. While the planktonic community was largely the same across depth, intraceullular metabolite pools were distinct in samples collected from the epilimnion vs. metalimnion and suggested depth-discrete partitioning of the arginine deiminase-mediated ornithine-arginine cycle (OAC, *i.e.,* urea cycle). Cumulatively, this study sheds light on the physiological processes that support *P. rubescens* blooms in the low-light metalimnion – suggesting the OAC plays a role in the ecological success of *P. rubescens* in this unusual niche. We subsequently demonstrate arginine deiminase (*argE*) is widely distributed across *Planktothrix* spp. and toxigenic cHAB genera broadly, implying *argE*-mediated arginine recycling may confer a competitive advantage to microcystin producers across a multitude of environmental stressors. This work lends support for the role of amino acids in determining microcystin congeners and inherent bloom toxicity – suggesting exogenous *vs.* endogenous supplies of arginine merit further attention. On a larger scale, this work strengthens the case that alternative nitrogen metabolism – including urea utilization, amino acid uptake, and the OAC – is a driver of toxigenic cyanobacterial blooms in fresh waters.

## Methods

### Meads Quarry field sampling

A bloom of reddish-pink, filamentous algae was initially noted in Meads Quarry (Knoxville, TN, USA) on March 13^th^, 2022 and April 13^th^, 2022, spurring the collection of preliminary microscopy and DNA samples (Supplemental Figure 1-3). Subsequently, a comprehensive field investigation was conducted by students on March 5^th^, 2024 at the field site (max depth = 7 m) (35°57’9” N, 83°51’57” W), where we again visually observed a reddish-pink bloom (Figure 1). Depth-discrete physiochemical profiles (Temperature, Conductivity, pH, Chlorophyll *a*, Phycocyanin, Turbidity) were documented using a pre-calibrated EXO2 Multiparameter Sonde (YSI Inc, Yellow Springs, OH, USA). The light climate was assessed at each depth using both a Secchi disk and a HOBO MX2202 light / temperature data logger (Onset, Bourne, MA, USA). Subsequently, three depths were selected for comprehensive sampling based on water column structure: 0.5 m (epilimnion), 2.0 m (metalimnion), and 6.0 m (near bottom) (Supplemental Table 1A). Samples were collected (n = 4) at each of the three depths with a Van Dorn bottle (4 L). For each replicate, 15 mL volumes of whole water were preserved with glutaraldehyde (2% final concentration) and stored in the dark at 4° C for assessment *via* FlowCAM analysis (Yokogawa Fluid Imaging Technologies, Scarborough, ME, USA). To collect samples for chlorophyll *a* (Chl *a*), 100 mL volumes of whole water were filtered through 0.22-μm (total Chl *a*) or 20.0-μm (large-size fraction Chl *a*) polycarbonate filters and stored at -20° C. Photopigment samples were collected by filtering 100 mL whole water samples onto GF-75 filters (Advantec MFS, Inc., Dublin, CA, USA) and stored at -80° C until processing. For toxin quantification (microcystins; 22 congeners, anatoxins; 6 congeners, cylindrospermopsins; 3 congeners), 100 mL volumes of whole water were collected onto GF-75 filters, flash frozen in liquid N_2_ and stored at -80°C until processing. For DNA analyses, whole water samples (180-600 mL, dependent of plankton density) were filtered through 0.22-μm pore-size Sterivex™, flash frozen in liquid nitrogen, and stored at -80° C until processing. Dissolved nutrient samples (50 mL volumes of Sterivex™ filtrate) were stored at -20° C until processing at the *The Ohio State University Stone Laboratory* on a QuAAtro 5-Channel continuous segmented flow auto-analyzer (Seal Analytical Inc., Mequon, WI, USA). For metabolomics, 120 mL volumes of whole water were filtered through 0.22 μm pore-size polycarbonate filters, flash frozen and stored at -80° C.

**Figure 1:**
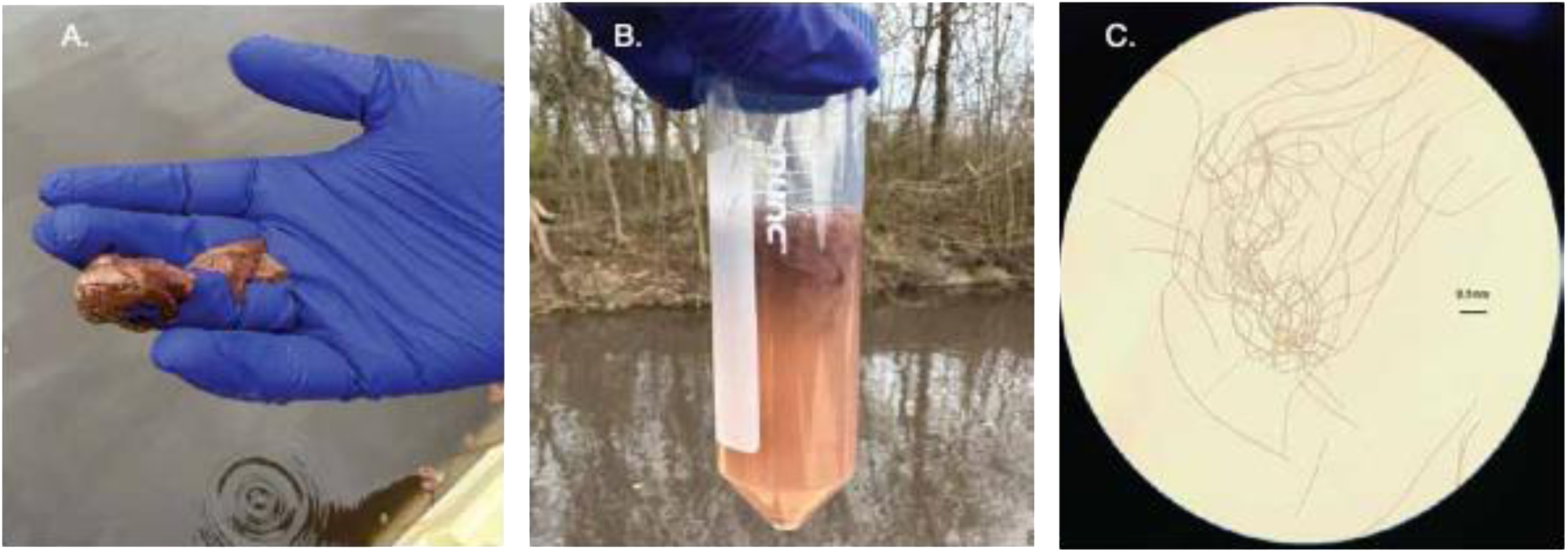
Qualitative observations of the *P. rubescens* bloom present in Meads Quarry (March 5^th^, 2024). (A) Pink algal surface scum found along the floating dock field site. Photo credit: Alexander R. Truchon. (B) Whole water sample demonstrating a dense, filamentous pink algal bloom. Photo credit: Brittany N. Zepernick. Brightfield microscopy image of whole water samples (100x magnification, wet mount) illustrating the presence of pink filaments. Photo credit: Jason L. Olavesen.

### Morphological analysis

FlowCAM 8000 imaging was performed using the 10x objective to generate morphological characterizations of algae in samples (Supplemental Table 1B). Briefly, ∼1,000 filaments were manually selected per sample (n = 4 biological replicates per depth) and analyzed for cylindrical biovolume (μm^3^) and length (μm).

Chl *a* samples were extracted in 90% acetone for 24 h at 4° C and quantified on a Turner Designs 10-AU Field Fluorometer (San Jose, CA, USA) to determine net concentrations (μg L ^-1^) (Welschmeyer, 1994). To characterize photopigments, three replicates collected from the metalimnion (depth = 2.0 m) were extracted in 100% acetone and subsequently incubated at -20° C for 24 has described previously (Paerl et al., 2014). Extracts were processed *via* high-performance liquid chromatography (HPLC) analysis at the *Institute of Marine Sciences - University of North Carolina at Chapel Hill* (Pinckney et al., 1996; Pinckney et al., 2001; Van Heukelem et al., 1994). Carotenoid identification was based on absorption spectra relative to purified pigment standards (DHI, Hørsholm, Denmark) with pigment concentrations reported in μg L ^-1^.

### Toxin extractions and assessment

Cyanotoxin samples (n = 12) were extracted in a 50% acidified methanolic solvent prior to analysis *via* HPLC-coupled with single quadruple mass spectrometry at the *State University of New York College of Environmental Science and Forestry*. Toxin congener identification was based on absorption spectra of primary standards (Enzo Life Sciences, Farmingdale, NY, USA) with intracellular toxin concentrations reported in μg L ^-1^. Refer to the published protocol for comprehensive details (Boyer, 2020).

### DNA extractions

Nucleic acid extractions were performed using standard basic-phenol extraction and ethanol precipitation as described previously (Martin and Wilhelm, 2020; Zepernick et al., 2022). DNA quality was assessed *via* Nanodrop ND-100 Spectrophotometry (Thermo Fisher Scientific, Waltham, MA, USA) and DNA quantity was determined using the Qubit dsDNA HS assay (Invitrogen, Waltham, MA, USA). Subsequently, two replicates from each depth were selected for sequencing along with the March and April 2022 samples (n = 8 total). Library preparation and sequencing were performed at *SeqCenter* (Pittsburg, PA, USA). Libraries were prepared using the Illumina DNA Prep kit (Illumina, San Diego, CA, USA) with a target insert size of 280 bp. Sequencing was performed on the Illumina NovaSeq X Plus generating ∼6 million 150 bp, paired-end reads for 2022 samples (n = 2) and ∼30 million 150 bp, paired-end reads for 2024 samples.

### Metagenomic analysis

Raw libraries (n = 8) were checked for quality *via* FastQC (v.0.12.1). Low quality reads were removed and adapters subsequently trimmed using Trimmomatic (leading = 20, trailing = 20, sliding window = 4:15, minimum length = 36) (v.0.39) (Bolger et al., 2014). Libraries were concatenated and assembled (coassembled) using MEGAHIT (v.1.2.9) (Li et al., 2015). Coassembly quality was assessed *via* QUAST (v.5.2.0) (Gurevich et al., 2013) and gene predictions (*i.e.,* open reading frames; ORFs) were called *via* MetaGeneMark (v.3.38) (Zhu et al., 2010). Prokaryotic taxonomic annotations were performed using ORF nucleic acid sequences and Kraken2 (v.2.1.3) (Kraken2 database = standard) (Wood et al., 2019). Functional annotations of ORF protein sequences were accomplished *via* eggNOG (v.2.1.12) (reference database = standard (v.5.0.2), e-value < 1e^-10^) (Cantalapiedra et al., 2021). Paired reads in each library were interleaved *via* reformat.sh available in BBTools (v.38.18) (Bushnell, 2014) then mapped to the coassembly using bbmap.sh. Following, tabulation of read counts to ORFs was performed *via* featureCounts in the subread package (v.2.0.2) (Bushnell, 2014). Normalization of read mappings to the coassembly (reads per kilobase per million; RPKM) (Dick, 2018) was accomplished in the ISAAC NG OnDemand RStudio Server (v.4.0.4). KEGG Orthology (Kanehisa et al., 2016) gene set enrichment was performed with all putative *Plankothrix* spp. mappings across time and depth using the R package clusterProfiler (v.4.10.1) (Settings: pAdjustMethod = Benjamini & Hochberg, minGSSize = 10, seed = 123) (Yu et al., 2012).

### Phylogenetic analysis

A phylogenetic tree was compiled to resolve species-specific challenges in the taxonomic annotations between *P. agardhii* and *P. rubescens,* given the suggestion that these two species are ecotypes (Pancrace et al., 2017). A manual reference database was curated using the Genome Taxonomy Database (GTDB) taxonomy tree (v.R214) (Chaumeil et al., 2020). Briefly, putative *rpoB* nucleotide sequences were extracted from all annotated genomes in GTDB for query against the *rpoB* nucleotide sequence most represented in Meads Quarry metagenomic libraries. To construct a family tree, *rpoB* nucleotide sequences were extracted from each species in the *Microcoleaceae* family. Multiple sequence alignments were performed for genes of interest using MAFFT (v.7.490) (Katoh and Standley, 2013) and subsequently trimmed using trimAl (settings: default) (v.1.4. rev15) (Capella-Gutiérrez et al., 2009). Maximum likelihood phylogenies were constructed using IQ-TREE (v.2.2.0.3) (von Haeseler et al., 2014) with an automated model finder (SYM+I+I+R2) and a bootstrap value of 100. Phylogenies were visualized using iTOL (v.6.9) (Ciccarelli et al., 2006).

A cyanobacterial arginine deiminase tree was compiled to explore the prevalence of arginine deiminase (*argE*) in cyanobacterial genera broadly and known microcystin producers. A manual reference database was curated to include NCBI blastp hits of our study *argE* protein to cyanobacteria (taxid:1117), *Microcystis* (taxid: 1125), Aphanizomenonaceae (taxid:1892259), and the prior reported cyanobacterial protein sequences of *argE (model organism-Anabaena but since reclassified as Dolichospermum)* (Lee and Rhee, 2020) and bifunctional *argZ* (model organism-*Synechocystis)* (Zhang et al., 2018). To construct a family tree, the same pipeline was followed as above with maximum likelihood phylogenies constructed using an automated model finder (Q.plant+R4) and a bootstrap value of 1,000.

### Metabolite extractions and UHPLC-HRMS analysis

Metabolite samples were immersed in extraction solvent (40% methanol, 20% water, 40% acetonitrile, and 0.1 M formic acid) and incubated at -20° C for 20 min to facilitate metabolite release (Lu et al., 2010; Rabinowitz and Kimball, 2007). Samples were centrifuged (2,500 xg for 5 min at 4° C) with the supernatant subsequently dried with nitrogen gas. Samples were reconstituted in 300 µL of LCMS grade water, vortexed, and centrifuged (2,500 xg for 5 min at 4° C) to ensure homogeneity.

Untargeted Ultra-High Performance Liquid Chromatography-High Resolution Mass Spectrometry (UHPLC-HRMS) was performed at the *University of Tennessee Biological and Small Molecule Mass Spectrometry Core via* the UltiMate 3000 RS chromatograph (Dionex, Sunnyvale, CA, USA) coupled to an Exactive™ Plus Orbitrap mass spectrometer (Thermo Fisher Scientific, Waltham, MA, USA). Separation of metabolites was achieved using a Synergi Hydro RP column (Phenomenex, Torrance, CA, USA) with dimensions of 100 × 2.0 mm and a particle size of 2.5μm, targeting smaller metabolites. Reversed-phase chromatography with an ion-pairing reagent was employed for the mobile phase (mobile phase composition for metabolite elution consisted of 97% water, 3% methanol, 11 mM tributylamine, and 15 mM acetic acid). A solvent gradient elution method was utilized to separate metabolites (Bazurto et al., 2018) at a flow rate of 0.2 mL min^-1^ and temperature of 25° C for 25 min. Metabolites eluted from the column were ionized in the electrospray ionization chamber using the negative polarity mode probe. Mass analysis was conducted using the orbitrap mass analyzer with a resolution of 140,000 within a scan range of 72-1000 *m z^-1^*. An injection time of 100 milliseconds and automatic gain control of 3 × 10^6^ was applied (Bazurto et al., 2018).

### Untargeted metabolomics analysis

Raw Xcalibur files were converted to centroided mzML format using the msConvert (v.3.0) (Adusumilli and Mallick, 2017) package within Proteowizard (Kessner et al., 2008). Subsequently, metabolites were identified based on a standards library *via* the Metabolomic Analysis and Visualization Engine (MAVEN) (mass tolerance ± 5 ppm to standard library) (v.0.12.0) (Melamud et al., 2010). This identification was validated manually using the natural abundance of isotopes in the compound. MAVEN was used to generate pre-processed peak data tables. Peak intensities were normalized by dividing the raw peak areas by the volume of sample filtered for metabolite extraction. Statistical analysis was performed with Metaboanalyst (v.6.0) (Pang et al., 2024). Variance filtering of data was performed using interquartile range (IQR), followed by log transformation and pareto scaling. Partial Least Squares-Discriminant Analysis (PLS-DA) was used to visualize the metabolic profile differences by depth. Metabolites with a Variable Important Projection (VIP) score ≥ 1 were deemed significantly differentially abundant and subsequently included in enrichment analyses. Statistical analyses were considered significant if p ≤ 0.05 and pathway significance was established at a false discovery rate (FDR) ≤ 0.05 (Clark et al., 2024).

## Results

### A cold, light-limited, nitrate replete metalimnion in Meads Quarry

In March 2024, the water column of Meads Quarry exhibited a thermocline: temperatures were ∼14° C in the epilimnion (0.5 m depth) decreasing to 6° C near bottom (6.0 m depth) (Figure 2A) (Supplemental Table 1C). We define “cold” as water column temperatures < 15° C in accordance to a prior study (Reinl et al., 2023), though we recognize the term is relative. Light attenuated with depth, exhibiting ∼40 μmol photons m^-1^ s^-2^ at 0.5 m (epilimnion) and rapidly decreasing to < 1 μmol photons m^-1^ s^-2^ at 2 m (metalimnion) (Figure 2B) (Supplemental Table 1C). Secchi depth was recorded at ∼1.0 m, supporting a turbid, light-limited metalimnion (Supplemental Table 1C). The pH of the water column was slightly basic (pH 8.50) at 0.5 m and gradually declined to 7.85 near-bottom (Figure 2C) (Supplemental Table 1C). While dissolved reactive phosphorus (DRP) was below detectable concentrations (< 0.3 μM), dissolved silicate ([dSi] = 59.3 μM +/-4.5) and nitrite ([NO ^-1^] = 0.5 μM +/-0.0) were present at all depths and demonstrated little variation (Supplemental Table 1D). Ammonium concentrations were highest near bottom ([NH_4_ ^+^] = 3.8 +/-0.4) and undetected at 0.5 and 2.0 m depths (Figure 2D) (Supplemental Table 1D). Nitrate concentrations were replete throughout the water column ([NO_3_] = > 17.0 μM +/-1.5) (Figure 2D) (Supplemental Table 1D).

**Figure 2:**
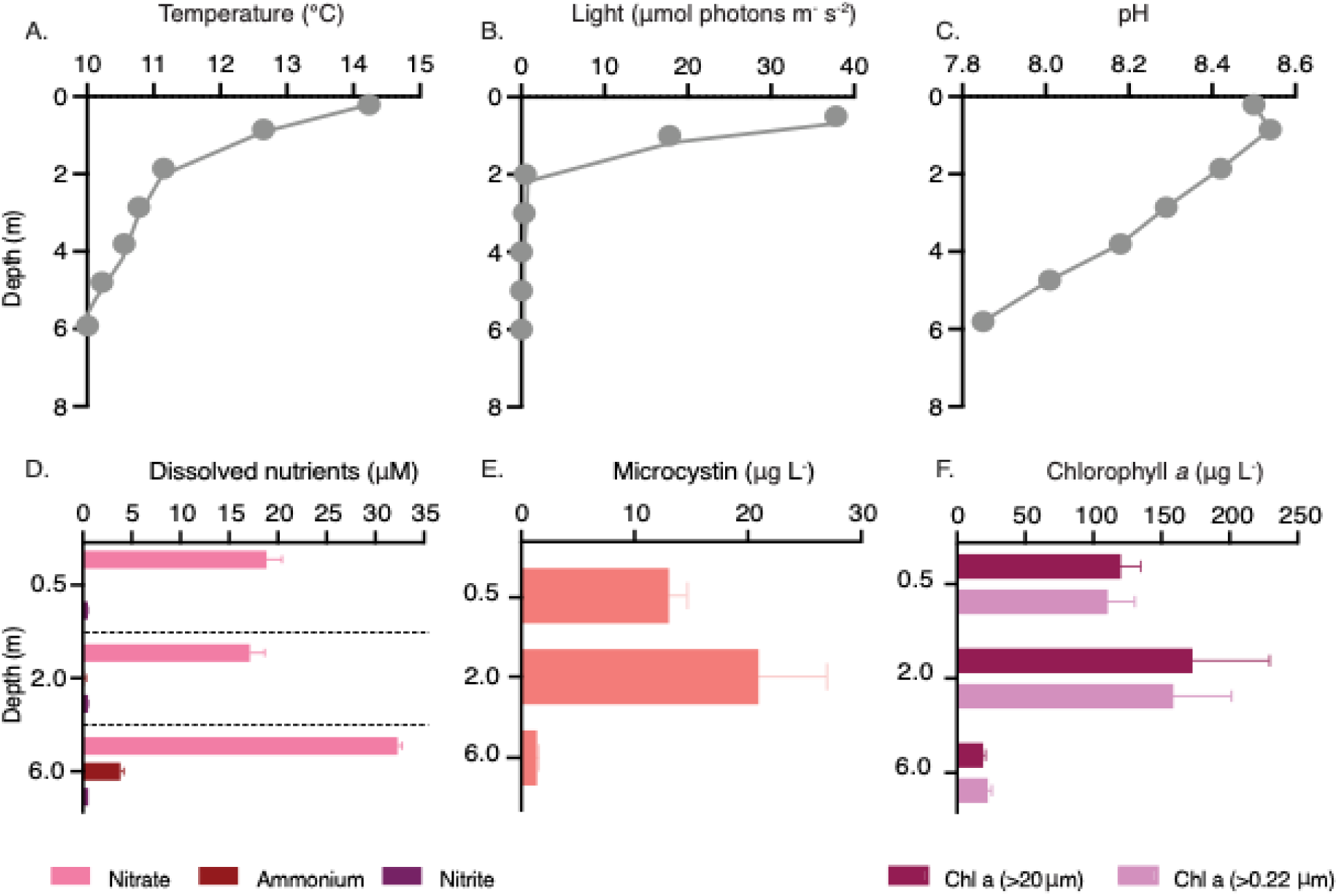
Physiochemical and biotic characterization of the Meads Quarry vertical water column. (A) Water temperature (°C) spanning the vertical profile of the water column at depths 0.5, 1.0, 2.0, 3.0, 4.0, 5.0 and 6.0 m. (B) Light intensity (μmol photons m^-1^ s^-2^) spanning the vertical profile of the water column. (C) pH spanning the vertical profile of the water column. (D) Dissolved nutrient concentrations of interest (μM) at select depths 0.5, 2.0 and 6.0 m. Nitrate concentrations are indicated in pink, ammonium concentrations are indicated in red, and nitrite concentrations are in purple. (E) Intracellular [DAsp^3^]MC-LR (μg L^-1^) in samples collected from 0.5, 2.0 and 6.0 m depths (salmon). (F) Chlorophyll *a* concentrations (μg L^-1^) of the total photosynthetic community (>0.22 μm size, light pink) and the large fraction of the photosynthetic community (>20 μm), dark pink). In panels D, E, and F, the mean (n = 4) is reported with the standard error of the mean (SEM) indicated by error bars.

**Figure 3:**
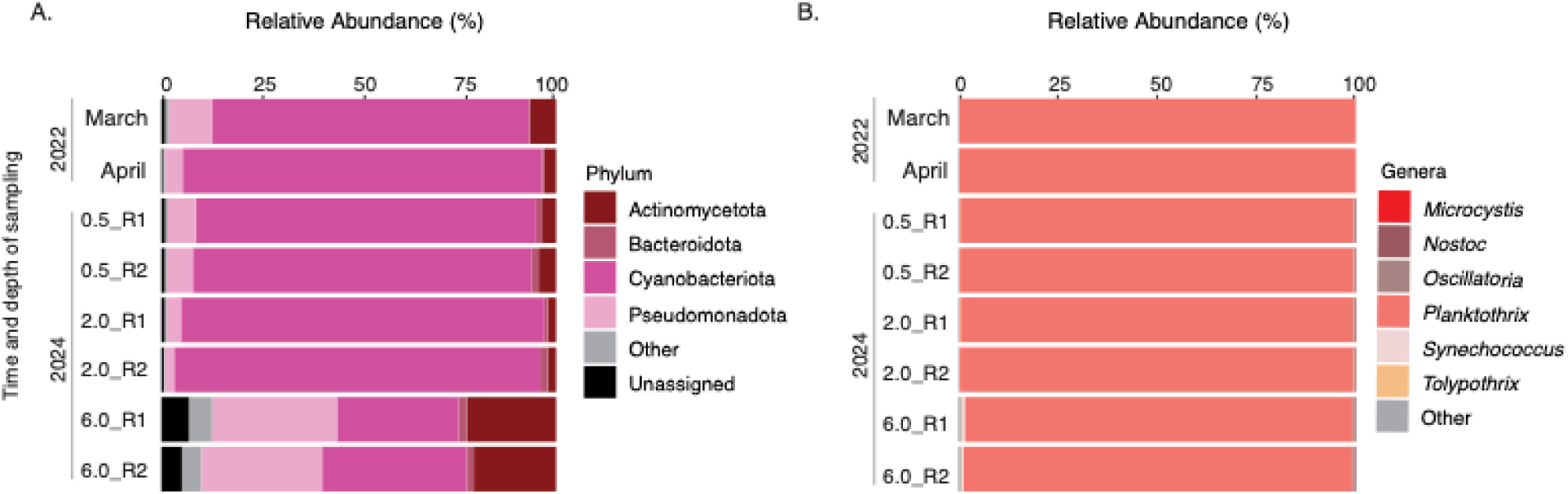
Relative abundance of mapped reads within each metagenomic library to coassembly. (A) Relative abundance of major prokaryotic phyla (%) across libraries. Opportunistic surface samples collected in March and April of 2022 are compared alongside the comprehensive vertical depth sampling conducted in March of 2024 (depths = 0.5, 2.0 and 6.0 m). Biological replication is indicated where applicable (R1 and R2). Actinomycetota = dark red, Bacteroidota = pale pink, Cyanobacteria = bright pink, Pseudomonadota = light pink, Other = grey, Unassigned = black. “Other” was determined as taxa contributing < 1% of total reads across all libraries (n = 39 phyla). (B) Relative abundance of major cyanobacterial genera (%) across libraries. *Microcystis* spp. = bright red, *Nostoc* spp. = mauve, *Oscillatoria* spp. = beige, *Planktothrix* spp. = salmon, *Synechococcus* spp. = pale pink, *Tolypothrix* spp. = peach, Other = grey. “Other” was determined as taxa contributing < 0.05% of total reads across all libraries (n = 55 assigned genera in addition to various unassigned genera).

### High concentrations of microcystin congeners detected throughout the water column

While anatoxins and cylindrospermopsins were below the limits of detection at all depths (<0.006 μg L^-1^ and <0.04 μg L^-1^, respectively), microcystin concentrations at 0.5 m and 2.0 m exceeded the EPA’s recommended threshold for recreational exposure ([microcystin] ≥ 8 μg L^-1^) (EPA, 2019). Microcystin concentrations were highest at 2.0 m (20.9 μg L^-1^ +/-6.1) and 0.5 m depth (13.0 μg L^-1^ +/-1.7) (Figure 2E) (Supplemental Table 1E), with ∼80% of microcystins within the epilimnion and metalimnion identified as congener [D-Asp^3^]MC-LR and ∼20% classified as the MC-YR congener. In addition, another prominent metbolite was identified with a target mass of *m/z* 1,024 – the mass matches that of six microcystin congeners. Prior reports investigating congeners synthesized by *P. rubescens* have reported the [D-Asp^3^]MC-RR congener at this target mass using NMR and MALDI-TOF mass spectrometry (Vasas et al., 2013). Yet, the metabolite in our study appeared to be distinct from the [D-Asp^3^]MC-RR congener standard and all microcystin standards tested (n = 22 microcystin congeners).

Specifically, this peak chromatographed several minutes earlier than [D-Asp^3^]MC-RR, and we could not resolve its UV spectrum at 239 nm. Subsequent identifications based on the molecular weight using CyanoMetDB (v. 03) (Jones et al., 2021) suggest the compound may alternatively be the cyanopeptide symplostatin 9 – a potent elastase inhibitor (PubChem CID #1: 71562563) (Salvador et al., 2013). Yet, lacking standards for this compound, this metabolite remains unidentified in our study.

### P. rubescens dominated the cyanobacterial community across the epilimnion and metalimnion

Cyanobacteria (estimated *via* phycocyanin; μg L^-1^) dominated the photosynthetic community (estimated *via* Chl *a*; μg L^-1^) (Supplemental Figure 4) (Supplemental Table 1C). Across both size fractions (Total Chl *a* > 0.22μm; Large size fraction > 20μm), Chl *a* concentrations were highest in the metalimnion, with the larger size fraction of the photosynthetic community demonstrating 172.5 μg L^-1^ +/-57.0 Chl *a* at 2.0 m depth, 119.9 μg L^-1^ +/-15.0 at 0.5 m, and 19.0 μg L^-1^ +/-1.9 at 6.0 m (Figure 2F) (Supplemental Table 1F).

FlowCAM analysis of ∼1,000 pink filaments per replicate revealed filaments were highest in biovolume (90,990.9 μm^3^ +/-3,532.1) and length (650.7 μm +/-23.8) at 2.0 m depth (Supplemental Figure 5) (Supplemental Table 1G). Accessory photopigment analyses indicated samples contained photopigments common to cyanobacteria, including myxoxanthophyll (13.6 μg L^-1^ +/-2.9), zeaxanthin (5.0 μg L^-1^ +/-1.0) and echinenone (2.9 μg L^-1^ +/-0.3) (Supplemental Figure 6) (Supplemental Table 1H).

Metagenomic read mappings to the coassembly revealed cyanobacteria dominated ≥ 80% of the prokaryotic community in March and April of 2022 (Figure 3A) (Supplemental Table 1I). Likewise, samples from 2024 indicated cyanobacteria were comparably abundant across the epilimnion and metalimnion, contributing 85-90% of prokaryotic reads. In contrast, cyanobacteria comprised just ∼30% of prokaryotic reads at 6.0 m depth. *Planktothrix* spp. contributed ≥ 97% of cyanobacterial reads across all libraries (Figure 3B) (Supplemental Table 1J), despite the presence of 55 distinct cyanobacterial genera detected at low abundances throughout the water column (Supplemental Table 1K). To validate these findings, relative abundance estimates were also calculated *via* the phyloseq R package (v.3.20) (McMurdie and Holmes, 2013), with samples yielding comparable results (Supplemental Figure 7).

Cumulatively, data suggest a *Planktothrix* spp. bloom dominated Meads Quarry in March-April of 2022 and March of 2024. Here, we define a bloom as one genus contributing > 55% of the photosynthetic community which alters the normal ecological function of the system (Zepernick et al., 2024).

Phylogenetic analysis *via* the most represented putative *rpoB* gene in our study (Meads *Planktothrix*) indicated this gene groups strongly with other putative *P. rubescens rpoB* sequences (Figure 4). We note the bloom-associated *rpoB* gene in our study shared the highest percent identity (99.9 %) with *P. agardhii* strain NIES-905, which was reclassified as *P. rubescens* in the GTDB release 214. Analysis at the family level (Microcoleaceae) indicated *Planktothrix* spp. *rpoB* sequences grouped closely with *Limnoraphis* and *Limnospira* genera (Supplemental Figure 8).

**Figure 4:**
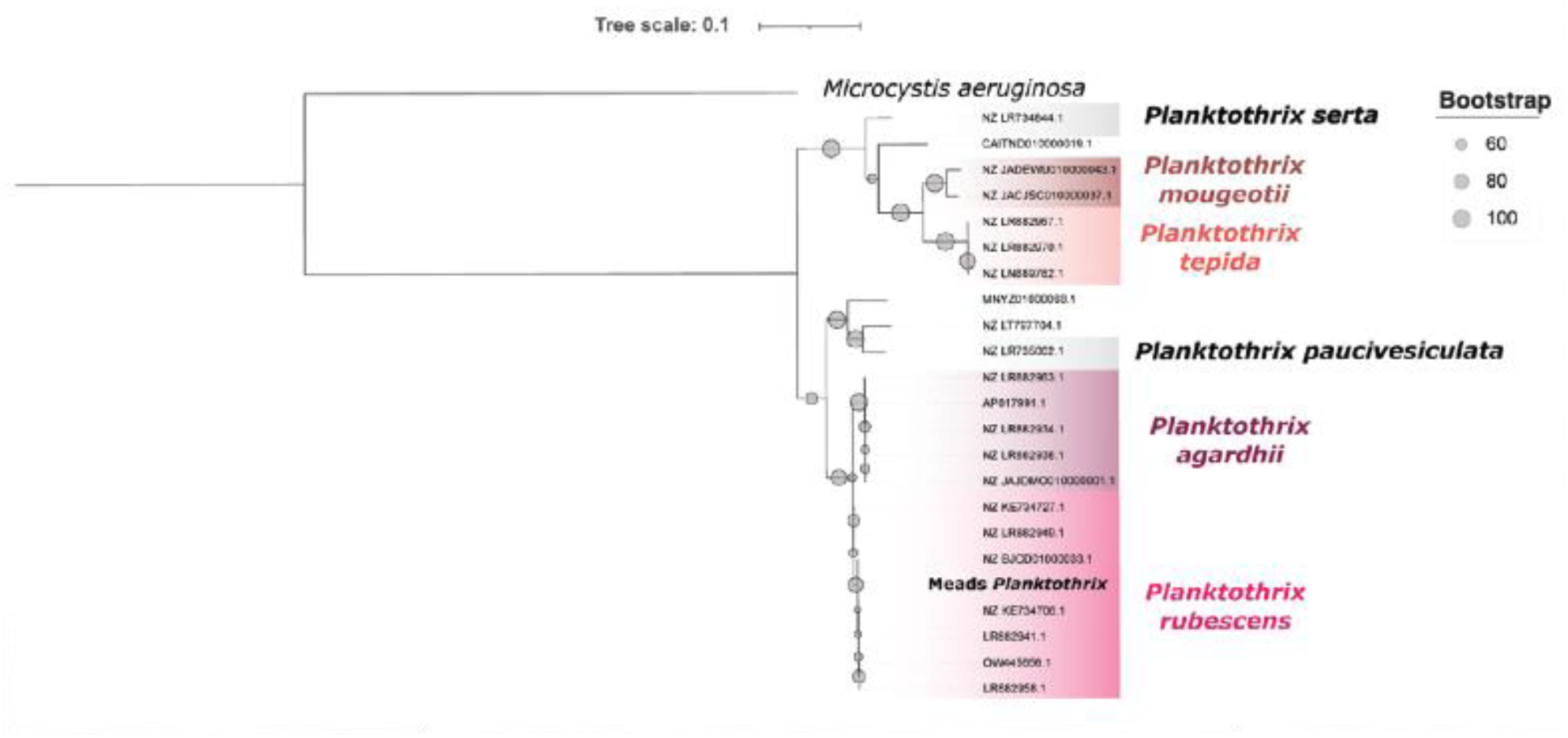
Phylogenetic reconstruction of the *Planktothrix* genus based on the nucleotide sequence of the *rpoB* gene. *Microcystis aeruginosa* was used as an outgroup for rooting. Bootstrap values are annotated at nodes by size. Defined species are annotated by color (*Planktothrix mougeotii* = mauve, *Planktothrix tepida* = salmon, *Planktothrix agardhii* = purple, *Planktothrix rubescens* = hot pink, Undefined species (*Planktothrix serta* and *paucivesiculata*) = grey).

### Nitrogen recycling and OAC genes enriched across the Planktothrix community

In total, 5,589 genes within the coassembly were annotated as putative *Planktothrix* genera – with the majority belonging to *P. rubescens* and a subpopulation annotated as *P. agardhii, P. tepida,* or unidentified at the species level (Supplemental Table 1L, M, N). These genes were statistically mined and leveraged with the literature to identify those which may have contributed to the ecological success of the Meads Quarry *Planktothrix* spp. community. KEGG enrichment of *Planktothrix* spp. gene abundances (RPKM) across all libraries (encompassing time and depth) indicated genes involved in amino acid biosynthesis (ko1230), amino sugar and nucleotide sugar metabolism (ko00520) and aminoacyl biosynthesis (ko00970) were significantly enriched (p_adj_ ≤0.05) (Figure 5A) (Supplemental Table 1O). Cumulatively, this suggests the *Planktothrix* spp. community within Meads Quarry possessed the genomic potential for nitrogen scavenging and recycling.

**Figure 5:**
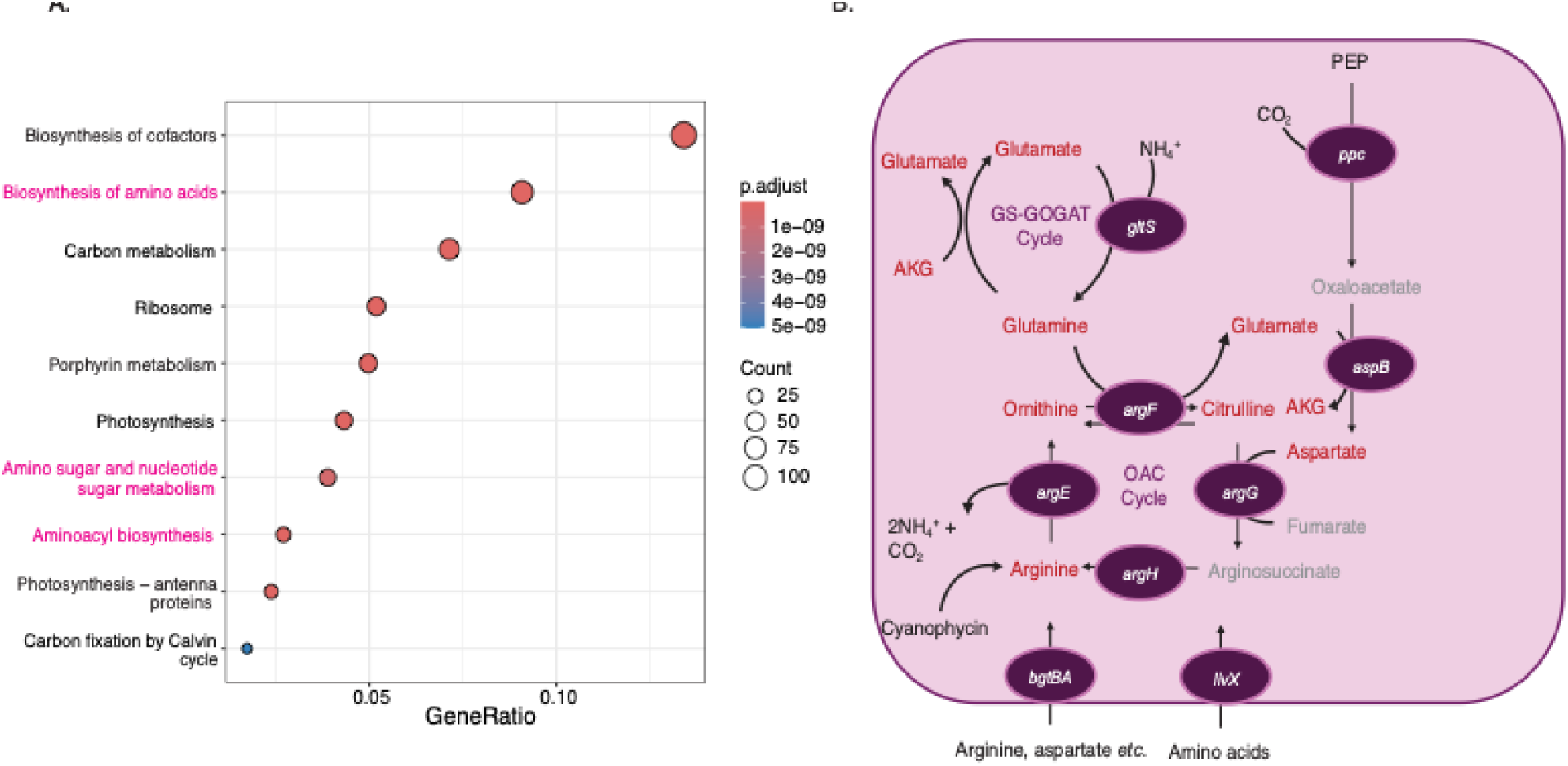
Analysis of putative *Planktothrix* spp. community gene enrichment and relative abundance. (A) Gene set enrichment analysis of all genes taxonomically annotated as *Planktothrix* spp. (n = 5,589) across all depths and time. Results are categorized by KEGG mapper metabolic pathways (n = 10) which are subsequently plotted by gene ratio and significance. Pathways of interest are indicated in hot pink font. (B) Schematic of the *argE*-mediated OAC pathway in cyanobacteria. Genes that were confirmed present in the Meads Quarry coassembly are indicated in dark pink circles. Metabolite reactants and products catalyzed by each gene are shown, with metabolites significantly enriched at 2.0 m depth in red font, those not detected in light grey font, and those detected but not significantly enriched at 2.0 m depth in black font.

In particular, two amino acid ABC transporters assembled: *livF,G,M,H* (K01996, K01995, K01998, K01997, gene_34483-34486) and *bztA,B* (K09969, K09970, gene_98217-98218). All genes involved in the OAC were recovered: arginine deiminase (*argE*; gene_28568), ornithine carbamoyltransferase (*argF*; K00611; gene_113736), argininosuccinate synthase (*argG*, KO1940; gene_28622) and arginosuccinate lyase (*argH*; K01755; gene_62500) (Figure 5B). Glutamate synthase of the GS-GOGAT cycle was recovered (*gltS*; K00284, gene_171375), along with phosphoenolpyruvate carboxylase (*ppc*; K01595; gene_106033) and aspartate aminotransferase (*aspB*; K00812; gene_211018) (Figure 5B). Cyanophycin synthetase (*cphA*) and cyanophycinase (*cphB*) were widely abundant in the *Planktothrix* spp. community and adjacently assembled (Supplemental Figure 9A) (Supplemental Table 1N), along with portions of microcystin synthetase (Supplemental Figure 9B) (Supplemental Table 1N). Urea transporters (*urtA,B,C,D,E*; K11959-K11963; gene_30833-30837) also assembled in addition to portions of urease (Supplemental Table 1N). In total, all the aforementioned genes are directly or indirectly involved in nitrogen scavenging and metabolism.

### OAC metabolites involved in nitrogen recycling were differently enriched across depth

Untargeted metabolomic analysis detected 74 metabolites across the vertical depth profile (Supplemental Figure 10). Notably, 43% of detected metabolites were identified as nucleoside/nucleotide analogues or mono/di/tri phosphates and 32% were identified as amino acid derivatives or carbohydrates involved in nutrient metabolism. Metabolomic profiles distinctly clustered by depth (Figure 6A), suggesting differences in intracellular metabolite pools given depths 0.5 m and 2.0 m were both dominated in composition by *P. rubescens*. Subsequent pathway enrichment analyses were performed with metabolites statistically driving depth clustering (VIP scores > 1.0) (Figure 6B). Notably, metabolites involved in D-amino metabolism and arginine biosynthesis were only significantly enriched in 2.0 vs. 0.5 m depth comparisons. (FDR p-value ≤ 0.05). Specifically, nine metabolites involved in arginine biosynthesis and ten metabolites involved in pyrimidine metabolism were significantly increased in relative abundance at 2.0 m depth compared to 0.5 m (FDR p-value ≤ 0.05) (Supplemental Figure 11A, B). All metabolic products of the OAC were significantly enriched in 2.0 m depth samples except oxaloacetate, fumarate, and arginosuccinate (Figure 5B).

**Figure 6:**
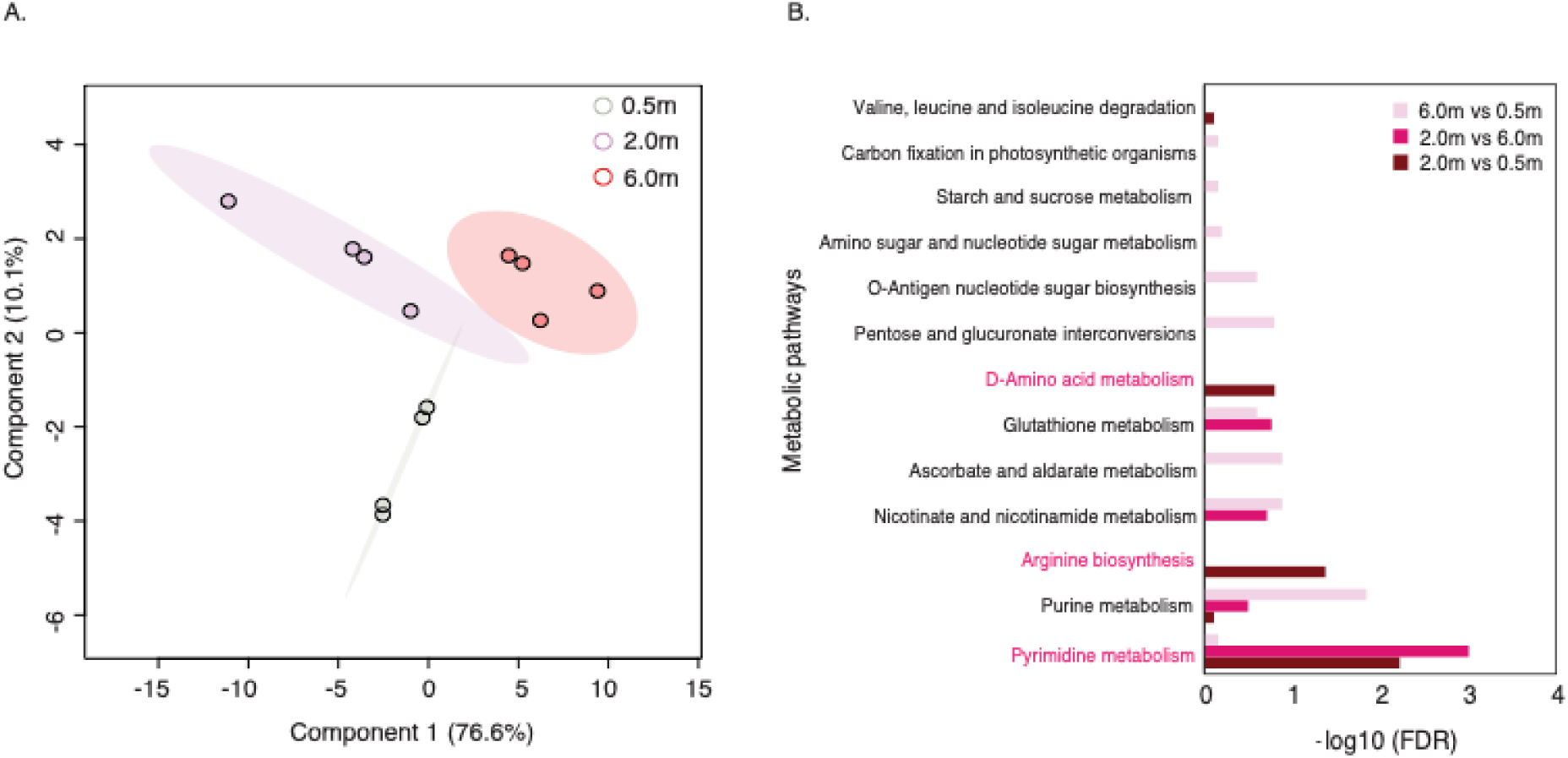
(A) Partial least squares discriminant analysis (PLS-DA) depicting clustering of metabolomic profiles (95% confidence interval) collected during March 2024 sampling efforts at 0.5 m (light grey), 2.0 m (light pink), and 6.0 m (salmon) depths. Each circle represents a biological replicate (n = 4 replicates per depth). (B) KEGG pathway enrichment analysis results by log-transformed FDR values for 6.0 vs. 0.5 m depth (light pink), 2.0 vs. 6.0 m depth (hot pink) and 2.0 vs. 0.5 m depth (dark red). Pathways of interest are indicated in hot pink font.

## Discussion

Conditions within the metalimnion routinely allow *P. rubescens* to outcompete other phytoplankton (Lenard, 2009; Walsby and Schanz, 2002; Zotina et al., 2003), with *P. rubescens’* affinity for the metalimnion suggested to serve as an evolutionarily stable strategy (Walsby et al., 2004). Studies have identified drivers contributing to the ecological success of *P. rubescens* in this niche (Selmeczy et al., 2025; Walsby and Schanz, 2002; Walsby et al., 2004) – with a persistent theme of the oligotrophic metalimnion (Ernst et al., 2009; Jacquet et al., 2005; Suarez et al., 2023). Yet, it remains to be mechanistically determined how *P. rubescens* routinely proliferates in the cold, low-light and often nutrient-limited metalimnion – at times on a perennial basis (Salmaso, 2000). Our results serve as a case study to hypothesize metabolic pathways which may facilitate the success of metalimnic *P. rubescens* blooms.

### Is a perennial P. rubescens bloom vertically cycling within Meads Quarry?

This study captured two ephemeral, epilimic blooms of *P. rubescens* during spring 2022 and 2024, with the latter spanning the epi-and metalimnion. Though low-light adapted *P. rubescens* typically exhibits growth and accumulation in the metalimnion (Bright and Walsby, 2000; Micheletti et al., 1998; Walsby and Schanz, 2002), specific conditions can induce the migration of metalimnic *P. rubescens* blooms to the epilimnion. This occasional event has been documented as the “*Burgundy-blood phenomenon*” – caused by low-light induced increases in cellular buoyancy coinciding with deep lake mixing and calm weather (Walsby et al., 2006).

While buoyancy regulation in cyanobacteria isn’t new, and prior studies have determined various cyanobacteria regulate their buoyancy in response to light and mixing regimes (Oliver and Walsby, 1984; Thomas and Walsby, 1985), *P. rubescens* contrasts against cHABs such as *Microcystis* spp., which migrate within the water column on a diel basis (Cui et al., 2016; Davenport et al., 2019; Hunter et al., 2008).

*P. rubescens* vertically migrates on a seasonal scale: populations accumulate within the metalimnion when high irradiances coincide with summer stratification, and subsequently migrate to the epilimnion when low irradiances coincide with deep-mixing and calm weather typical in fall / spring turnover (Lenard, 2009; Walsby et al., 2004; Walsby et al., 2006). We hypothesize our March-April 2022 and March 2024 sampling efforts captured epilimic *P. rubescens* blooms spurred by deep mixing throughout an isothermal winter-spring column. Further, our data suggest these recurring blooms in 2022 and 2024 may have been a result of annually dominant, metalimnic populations which reseed the epilimnion – implying *P. rubescens* may proliferate year-round in the quarry as documented in other northern, temperate lakes (Jacquet et al., 2005; Van den Wyngaert et al., 2011; Yankova et al., 2016). Indeed, Lake Zurich, Germany (Walsby et al., 2006), Lake Bourget, France (Jacquet et al., 2005), and Lake Mondsee, Austria (Dokulil and Teubner, 2012) experience seasonally persistent, long-term metalimnic *P. rubescens* blooms distinct from seasonal succession patterns of other cHABs – which bloom in the summer and overwinter in the sediments (Legnani et al., 2005; Moiron et al., 2021; Van den Wyngaert et al., 2011). Yet, little research is available concerning the annual variability of Meads Quarry’s seasonal profile – as is common in smaller lakes across the globe (Downing, 2010). Hence, small-scale systems such as Meads Quarry represent potential hotspots for undetected, seasonally-persistent *P. rubescens* blooms.

### Evidence the OAC plays an enhanced role in maintaining metalimnic P. rubescens blooms

Metabolites involved in amino acid metabolism and the OAC were significantly enriched in metalimnic samples – implying nitrogen scavenging and recycling in *P. rubescens* filaments. Though NO ^-^ was replete across depth and *P. rubescens* possessed the genomic potential to assimilate it (*i.e.,* NO_3_ ^-^ transporters), we hypothesize *P. rubescens* preferentially assimilated NH_4_ ^+^ as a primary nitrogen source which depleted concentrations across the epilimnion and metalimnion. This hypothesis is based upon prior observations by Atwood (2022), who noted NO_3_ ^-^ concentrations increased over the seasonal progression of an epilimnic *P. rubescens* bloom (February-March, Skinn Quarry, OH, USA). In turn, Jacquet et al. (2005) noted NO_3_ ^-^ concentrations were depleted above, not within, a metalimnic *P. rubescens* bloom (June-September, Lac du Bourget, France). Yet, this finding would only exacerbate the typically nutrient-poor nature of the metalimnion – and it fails to address how *P. rubescens* proliferated in this study.

The metalimnion is often light-limited compared to the light-replete epilimnion. Thus, carbon fixation and ATP synthesis *via* photosynthesis are coincidingly decreased. Oligotrophic conditions and biotic drawdown of nutrients compound this competitive conundrum in a stratified environment – raising the question how *P. rubescens* proliferates in this niche (Walsby et al., 2004). The data from our *P. rubescens*-dominated metabolomes distinctly clustered by depth – with OAC (*i.e.,* urea cycle) metabolite pools driving this disparity *via* significant enrichment in metalimnic filaments. The OAC has been well-characterized in cyanobacteria, with studies demonstrating *Microcysti*s spp. uses urea as a carbon and nitrogen source to evade pH-induced carbon and nitrogen limitation during epilimnic, summer blooms (Belisle et al., 2016; Krausfeldt et al., 2019; Steffen et al., 2017). We build off these studies by suggesting *P. rubescens* uses the OAC to evade carbon and nitrogen limitation in the low-light metalimnion. In further support, recent studies identified arginine deiminase (*argZ*; *Synechocystis* spp. / *argE*; *Dolichospermum* spp.) as the rate-limiting step of the OAC in cyanobacteria – noting this enzyme cyclically converts arginine to ornithine (yielding 2 NH_4_ ^+^ + CO_2_ per cycle) (Burnat et al., 2019; Lee and Rhee, 2020; Zhang et al., 2018). These studies revealed the OAC not only allows cyanobacteria to use different nitrogen and carbon sources (*i.e.,* urea or amino acids), but also rapidly recycle these products – specifically arginine (Zhang et al., 2018) (Figure 5B). The question remains as to what compounds are assimilated into the OAC, how this is energetically driven and to what extent light climate constrains this cycle within the low-light metalimnion.

Sustained intracellular recycling by the *argE*-mediated OAC requires an influx of arginine. Though uptake assays were not performed in this study, we offer hypotheses concerning the source of arginine leveraged with the prior literature. The first logical hypothesis is the uptake, metabolism and subsequent integration of urea into the OAC (Krausfeldt et al., 2019; Steffen et al., 2017). While this remains probable, a prior study recorded negligible uptake rates of urea under low irradiances (3.5-11μmol photons m^-1^ s^-2^) and darkness in *P. rubescens* (Zotina et al., 2003) – suggesting this may not be a preferred source under light-limited and energetically constrained conditions. Alternatively, the uptake, metabolism and subsequent integration of amino acids into the OAC serves as an alternative source of arginine. This hypothesis is based upon observations describing uptake and assimilation of various amino acids by *P. rubescens* (Walsby and Jüttner, 2006; Zotina et al., 2003), with the light saturation irradiance for this process reported at 1 μmol photons m^-1^ s^-2^ (Walsby and Jüttner, 2006). While many amino acids also require metabolism before shuttling into the OAC, direct uptake of environmental arginine would circumvent energetic strain on this process – with a prior study reporting high uptake rates of arginine under low irradiances (3.5-11μmol photons m^-1^ s^-2^) and darkness in *P. rubescens* (Zotina et al., 2003). Lastly, arginine could be sourced intracellularly from cyanophycin (Flores et al., 2019) although the depletion of internal nitrogen reserves is not a long-term, sustainable mechanism for a cHAB that spends months in the low-light metalimnion. In summary, we suggest the *argE*-mediated OAC may be of enhanced metabolic importance in *P. rubescens* blooms within the light and nutrient limited metalimnion, but further research is required concerning the source of arginine influx and environmental constraints.

### A broader role of argE and amino acid availability in toxigenic cyanobacterial bloom ecology?

In cyanobacteria, arginine deiminase (*argE* / Z) was recently found to play a substantial role in the OAC and cellular responses to oscillating nitrogen conditions (Burnat et al., 2019; Lee and Rhee, 2020; Zhang et al., 2018). To date, *argE* has been characterized within the freshwater cHAB genera *Dolichospermum* spp. (Burnat et al., 2019; Lee and Rhee, 2020) – a notable microcystin producer which routinely blooms in the warm, light-replete epilimnion (Huisman et al., 2018). We determined *argE* is widely conserved within *Planktothrix* spp. and toxigenic cyanobacterial genera broadly (Figure 7). Notably, *argE* sequences distinctly clustered by genera – with *argE* recovered in a variety of known microcystin producers including *Planktothrix, Lyngbya, Microcoleus, Microcystis, Fischerella, Nostoc, Cylindrospermopsis (*recently renamed *Raphidioposis),* and *Dolichospermum* spp. (Figure 7). Cumulatively, we suggest *argE*-mediated arginine recycling *via* the OAC may shape the ecological success of toxigenic cHAB genera – which require ample nitrogen to synthesize arginine-rich microcystins.

**Figure 7:**
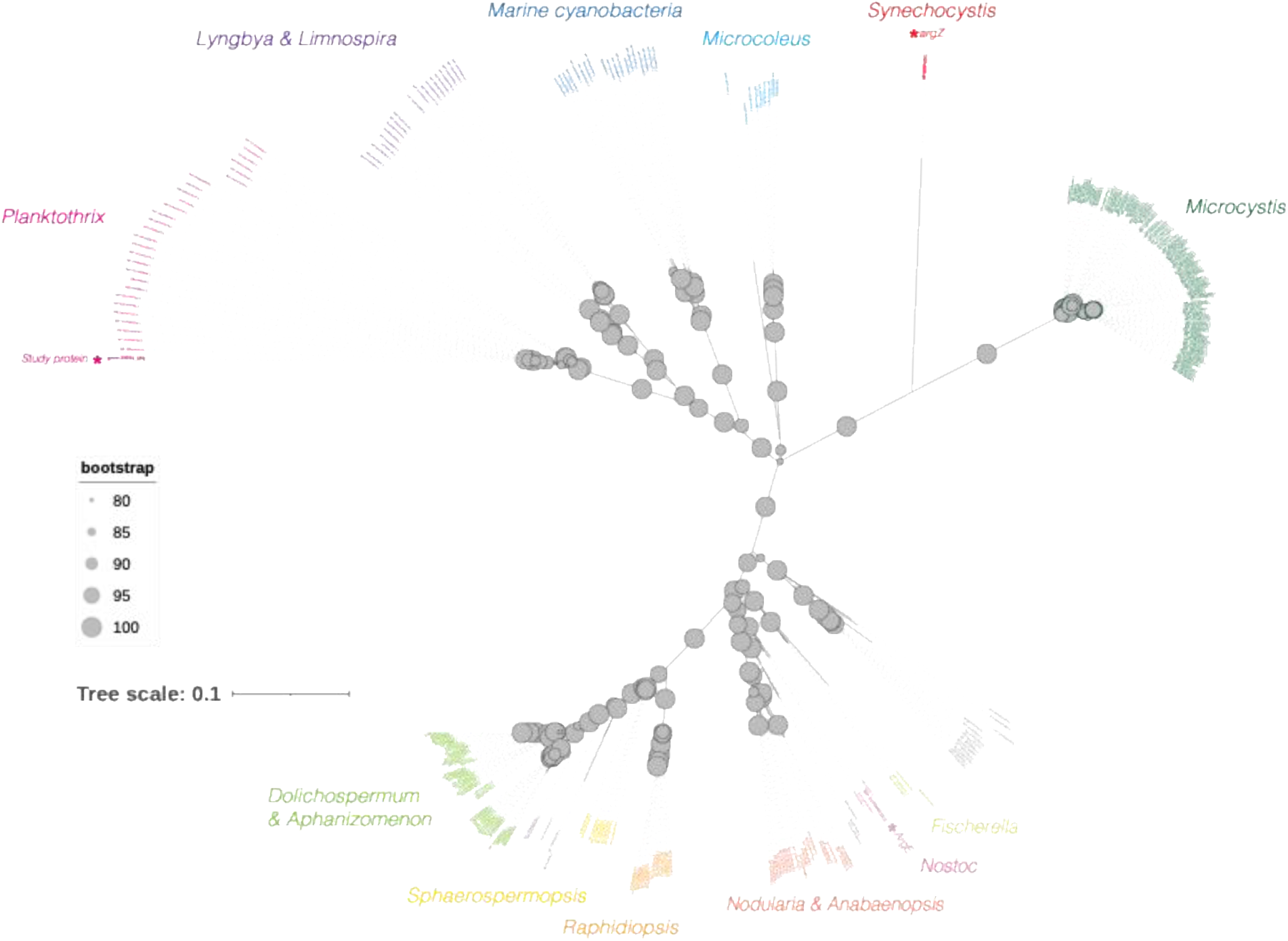
Phylogenetic reconstruction of putative *argE* protein sequences with respect to cyanobacteria. The *P. rubescens argE* protein in this study is denoted with an asterisk as “study protein” within the *Planktothrix* cluster (hot pink). The *argZ p*rotein sequence described within *Synechocystis* by Zhang et al. (2018) is indicated with an asterisk in red. The *argE* protein sequence described within *Anabaena* (since reclassified as *Dolichospermum*) by Lee and Rhee (2020) is indicated with an asterisk within the *Nostoc* cluster (light pink). Additional conserved clusters of the *argE* protein are labeled as follows: *Lyngbya* and *Limnospira* (purple), Marine cyanobacteria (dark blue), *Microcoleus* (light blue), *Microcystis* (dark green), *Fischerella* (yellow-green), *Nodularia* and *Anabaenopsis* (salmon), *Raphidiopsis* (orange), *Sphaerospermopsis* (yellow), *Dolichospermum* and *Aphanizomenon* (Aphanizomenonaceae; light green). Bootstrap values are in light grey, solid circles with size indicative of the value.

In turn, this suggests *argE*-mediated arginine recycling may serve an enhanced role in the fitness of microcystin producers beyond supplementing nitrogen metabolism. For example, arginine-rich microcystins have been suggested to confer fitness advantages in *Microcystis* spp. under oxidative stress and cold-temperature acclimation (Peng et al., 2018; Stark et al., 2023; Zilliges et al., 2011). With the epilimnic phenomenon in this study coinciding with cold temperatures and high irradiances, the ability to rapidly recycle arginine and synthesize microcystin would be particularly advantageous in this multi-stressor scenario. Indeed, microcystin concentrations were well above the level of recreational exposure in our study (EPA, 2019), with microcystin levels doubled in the metalimnion compared to the epilimnion.

Cumulatively, these results indicate further research is required regarding the role of *argE* and the OAC in cyanobacterial responses to environmental stressors – particularly at the interplay of light, temperature and nitrogen stress.

Beyond arginine recycling, this work underscores the role of amino acid availability in determining microcystin congeners. While the mechanisms which constrain microcystin variants in cyanobacteria remain debated, prior studies suggest microcystin congeners are largely determined by nitrogen species and availability (Chaffin et al., 2023; Monchamp et al., 2014; Puddick et al., 2016) – with exogenous and endogenous concentrations of arginine suggested to play a key role (Dai et al., 2019; Guo et al., 2023; Van de Waal et al., 2010). The more toxic microcystin-LR congener incorporates leucine at the first variable amino acid position, wheras the less-toxic microcystin-RR congener incorporates arginine. Hence, the availability of arginine is suggested to shape the dominant microcystin congener and inherent toxicity of the bloom. In support, a prior study noted exogenous arginine additions to *P. agardhii* monocultures decreased the microcystin LR/RR ratio (Tonk et al., 2008). In turn, another study noted additions of exogenous nitrate to *P. agardhii* monocultures decreased the [D-Asp^3^]MC LR/RR ratio – citing the increase in endogenous arginine *via* cyanophycin as the driver (Van de Waal et al., 2010). In contrast, we detected high concentrations of [D-Asp^3^]MC-LR in our environmental *P. rubescens* bloom and a mystery peak which possessed the same mass as [D-Asp^3^]MC-RR yet appeared to be distinct. While this trend is in accordance with a prior *P. agardhii* study that reported a decoupling of this phenomenon when extrapolated to the environment (Tonk et al., 2008) – it also suggests the factors which contstrain microcystin congener dominance are indeed complicated and multi-faceted. Cumulatively, this study builds the case for a deeper dive into how exogenous vs. endogenous arginine supply, amino acid availability, and nitrogen species broadly constrain microcystin congener biosynthesis (and thus toxcicity) in cyanobacterial genera.

## Conclusions

We leveraged a case study to generate ecophysiological hypotheses underlying the peculiar proliferation of metalimnetic *P. rubescens blooms* and built upon the case that alternative nitrogen metabolism – including urea utilization, amino acid uptake, and the OAC – is a driver of toxigenic cyanobacterial blooms in fresh waters. These findings contribute to historical studies describing the puzzling ecophysiology of *P. rubescens* and suggest *argE* (and the implicitly-related supply of exogenous / endogenous arginine) play a large role in the ecological success and toxicity of metalimnic *P. rubescens* blooms and microcystin-producers broadly. In turn, our study provides context to how this toxigenic cyanobacterium (and others possessing the *argE*-mediated OAC pathway) will fare in future scenarios. As climate change continues to exacerbate episodic mixing events and long-term stratification trends alike, there is a need to deduce how key players in lake ecosystems will respond. Here, we provide evidence that *P. rubescens* may serve as a formidable competitor in the increasingly stratitified (and thus nutrient and light limited) water column – particularly within deep, temperate resevoirs in northern latitudes. In tandem with our findings, this study presents a new array of questions at the interplay of nitrogen, cyanotoxins and a dynamic environment - indicating further research is required into the factors which determine microcystin congener synthesis.

## Data Availability

All raw metagenomic sequencing data generated by the *2024 MICR 669 Advanced Techniques in Field Microbiology* course are available on the NCBI Sequence Read Archive under BioProject PRJNA1162508, BioSamples: [SAMN43814365](https://www.ncbi.nlm.nih.gov/biosample/43814365), [SAMN43814366](https://www.ncbi.nlm.nih.gov/biosample/43814366), [SAMN43814367](https://www.ncbi.nlm.nih.gov/biosample/43814367), [SAMN43814368](https://www.ncbi.nlm.nih.gov/biosample/43814368), [SAMN43814369](https://www.ncbi.nlm.nih.gov/biosample/43814369), [SAMN43814370](https://www.ncbi.nlm.nih.gov/biosample/43814370), [SAMN43814371](https://www.ncbi.nlm.nih.gov/biosample/43814371), [SAMN43814372](https://www.ncbi.nlm.nih.gov/biosample/43814372), SRA: SRX26131638, SRX26131639, SRX26131640, SRX26131641, SRX26131642, SRX26131643, SRX26131644, SRX26131645 (Supplemental Table 1L). The coassembly generated from the 2022 and 2024 metagenomic libraries (n = 8) is available on [Zenodo](https://zenodo.org/records/13787465) (Zepernick, 2024). Metabolomics data are available under project PR002207 on the NIH Common Fund’s National Metabolomics Data Repository website the [Metabolomics Workbench](https://www.metabolomicsworkbench.org/) (Study ID ST003577). Data may be accessed directly *via* its assigned project doi: http://dx.doi.org/10.21228/M8BN6T. All environmental metadata are available in the Supplemental (Supplemental Tables 1B-H).

## Author Contributions

The data presented within this manuscript were collected as part of a collaborative, interdisciplinary effort between graduate students enrolled in *MICR 669 Advanced Techniques in Field Microbiology.* This course was designed and implemented by B.N.Z. who served as the primary instructor of record. Opportunistic samples collected from Meads Quarry in March and April of 2022 were collected by B.N.Z. The subsequent 2024 field survey of Meads Quarry was designed and implemented by graduate students enrolled in the course: B.A.A, M.J.A., K.A.H., J.L.O., Q. S., A.R.T., and J.L.W. DNA extractions were performed by B.N.Z., metabolite extractions were performed by B.A.A. and Q.S. Chl *a* extractions and FlowCAM analysis were performed by D.J.N., Photopigment analysis was performed by J.H.C. and H.W.P. Dissolved nutrient analysis was performed by K.S., J.D.C., and toxin analysis was performed by G.L.B. Physiochemical data analysis was performed by J.L.W., metagenomic analysis was performed by K.A.H., J.L.O., A.R.T., M.L.A. and B.N.Z. Metabolomic analysis was performed by B.A.A. Q.S. H.F.C and S.R.C. L.N.H aided in interpretation of metagenomic, metabolomic, and microcystin congener results. D.J.N. served as the Field Teaching Assistant and E.E.C. provided bioinformatic support as the Bioinformatic Teaching Assistant. Initial manuscript draft was written by B.N.Z. All authors contributed to the final version of the manuscript.

## Acknowledgements

We thank John D. Wehr at Fordham University for his expertise in phytoplankton identification, Katarina Jones for providing the Mass Spectrometry Core services and Joseph P. Caniglia for facilitating opportunistic collections of 2022 Meads Quarry samples. We also thank the investigators of the Great Lakes Center for Fresh Waters and Human Health (G.S.B., J.D.C., H.W.P., G.L.B., and S.W.W.) and the SEC Emerging Scholars Program for their collaboration and support in this project. We thank Ijams Nature Center’s Executive Director, Amber Parker, for her collaboration and support throughout this project. We note a portion of the computation for this work was performed on the *University of Tennessee Infrastructure for Scientific Applications and Advanced Computing (ISAAC)* computational resources.

