## Supplemental for "The ornithine-arginine cycle supported a toxic, metalimnic *Planktothrix rubescens* bloom"

For publication in conjunction with the following:

**Short title (100 characters max):** OAC maintained metalimnic *P. rubescens* bloom

**Key words:** Metagenomics, metabolomics, nitrogen metabolism, arginine deiminase, cold cyanobacterial bloom, microcystin congeners, urea

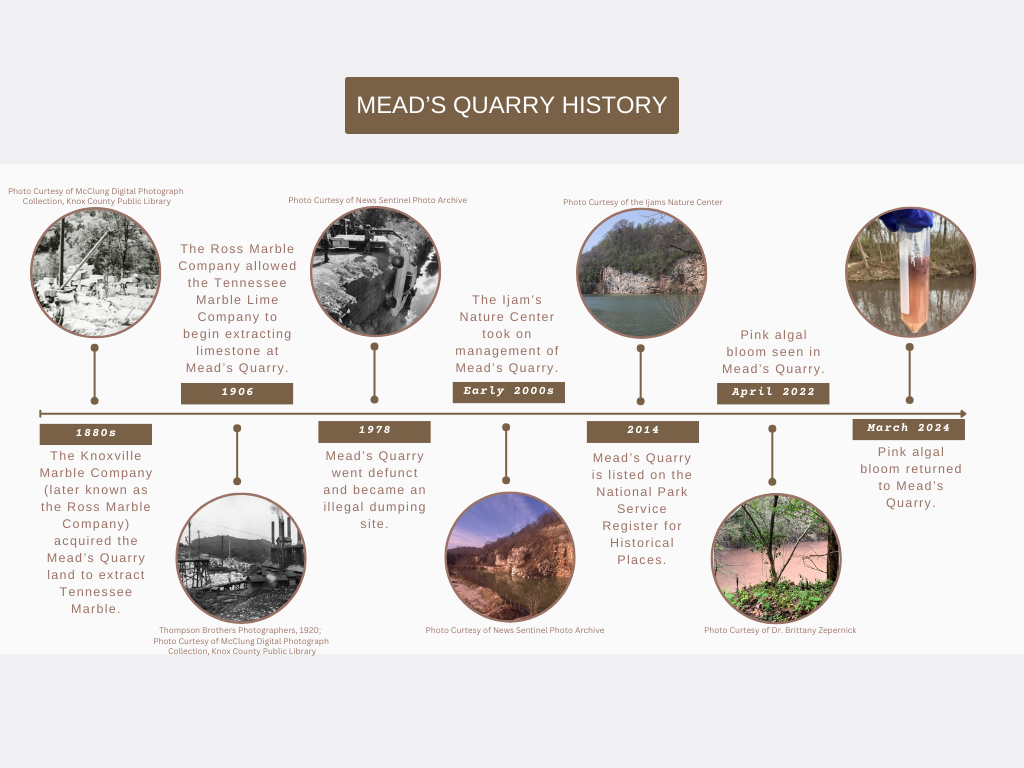

Supplemental Figure 1: Mead’s Quarry historical timeline leading up to algal bloom appearance. Photo sources are listed above or below each photo not owned by the authors. Photo credit: Jillian Walton.

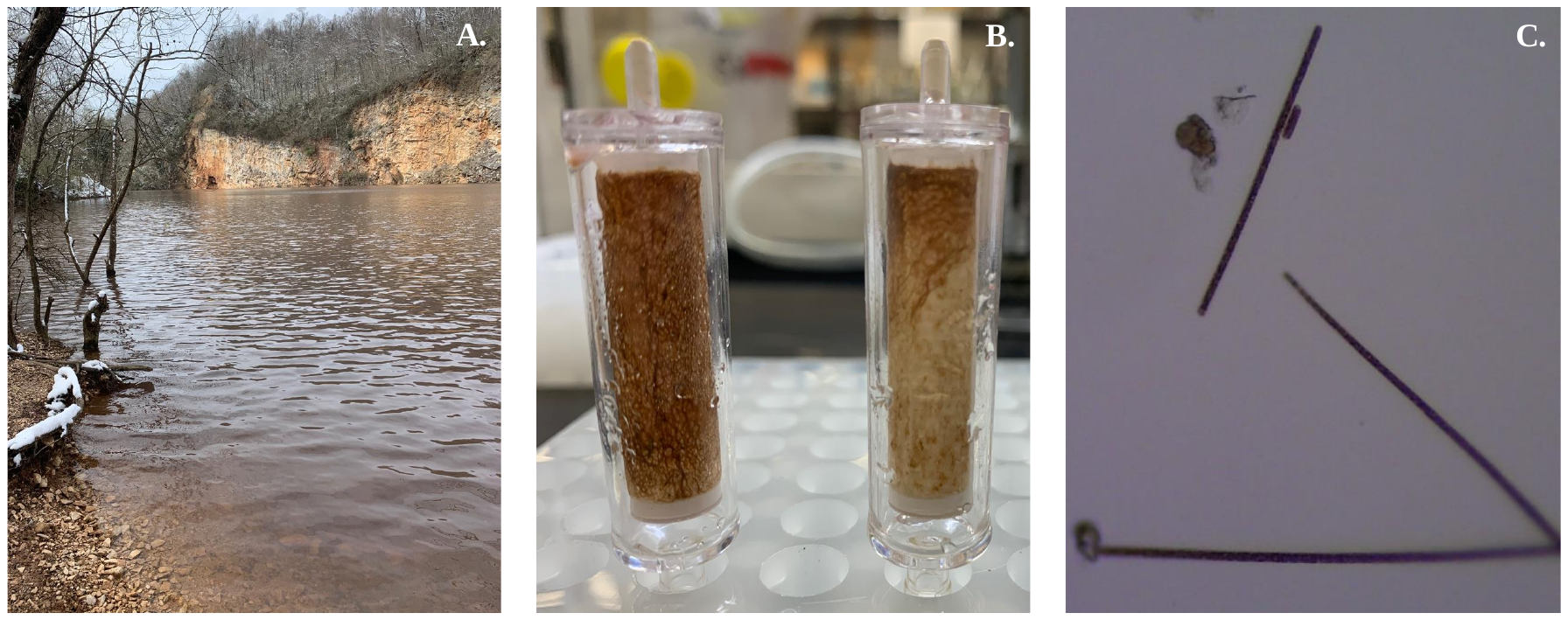

Supplemental Figure 2: Mystery pink, filamentous bloom at Mead’s Quarry found during a recreational visit on March 13^th^, 2022. (A) Image taken of the quarry before collecting opportunistic below-surface samples for subsequent DNA extraction. (B) Sterivex filters containing bright red/pink filaments. (C) Bright field microcopy image taken using a 40x objective. Photo credit: Brittany N. Zepernick.

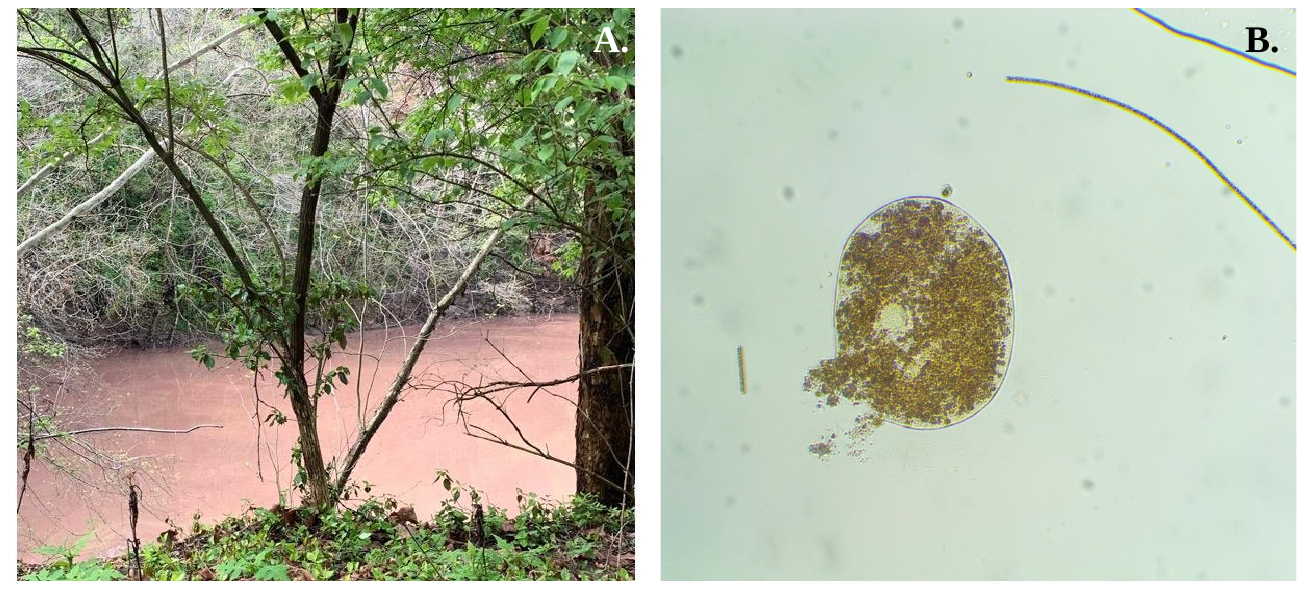

Supplemental Figure 3: Mystery pink, filamentous bloom at Meads Quarry found during a subsequent visit on April 13^th^, 2022. (A) Location where opportunistic sub-surface samples were collected for DNA extraction and microscopy. (B). Bright field microcopy image taken using a 10x objective. Photo credit: Brittany N. Zepernick.

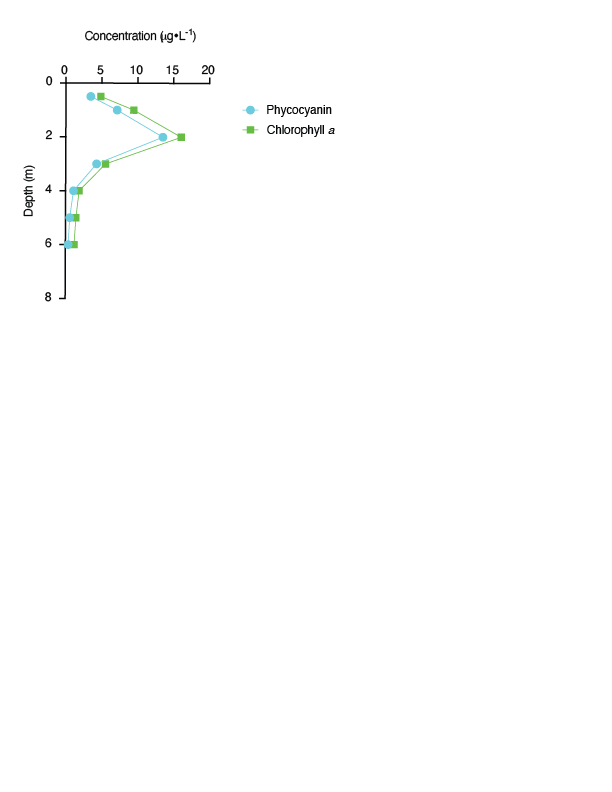

Supplemental Figure 4: EXO2 sonde metric at 0.5, 1.0, 2.0, 3.0, 4.0, 5.0 and 6.0 m depths. Phycocyanin concentrations representing the total cyanobacterial community are indicated in blue (μg L^-1^). Chlorophyll *a* concentrations representing the total photosynthetic community are indicated in green (μg L^-1^).

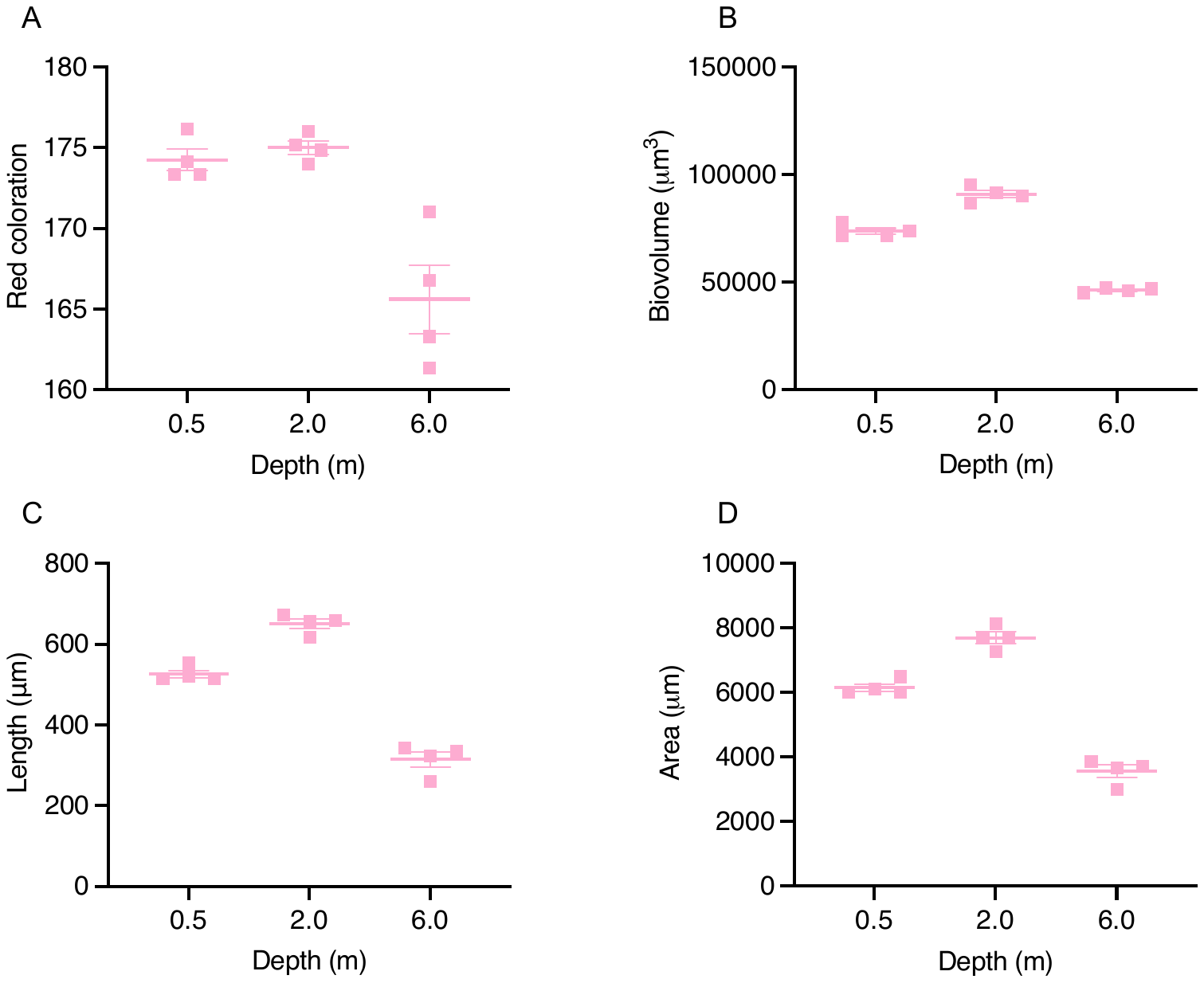

Supplemental Figure 5: FlowCAM morphological metrics of pink cyanobacterial filaments collected from the depths of 0.5, 2.0 and 6.0 m. Each square is the mean of the total filaments measured in that biological replicate (n = ~1,000), with the mean of replicates indicated by a horizontal line, and the standard error of the mean indicated with vertical error bars. (A) Average red color of filaments at each depth. (B) Average biovolume (μm^3^) of filaments at each depth. (C) Average length (μm) of filaments at each depth. (D) Average area of filaments at each depth.

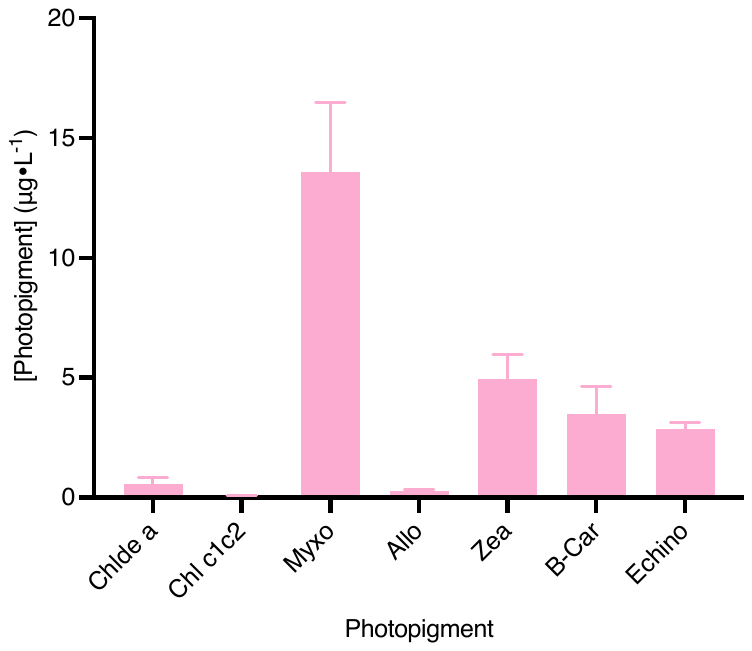

Supplemental Figure 6: Intracellular photopigment concentrations (μg L^-1^) of pink filaments collected from 2.0 m depth. Chlde *a* = chlorophyllide *a*, Chl c1c2 = chlorophyll *c_1_c_2_* Myxo = myxoxanthophyll, Allo = alloxanthin, Zea = zeaxanthin, B-car = β-carotene, Echino = echinenone. Only photopigments that were detected within the samples are shown. Photopigments that were not detected but screened for included fucoxanthin, neoxanthin, violaxanthin, diadinoxanthin, antheraxanthin, canthaxanthin and chlorophyll *b*.

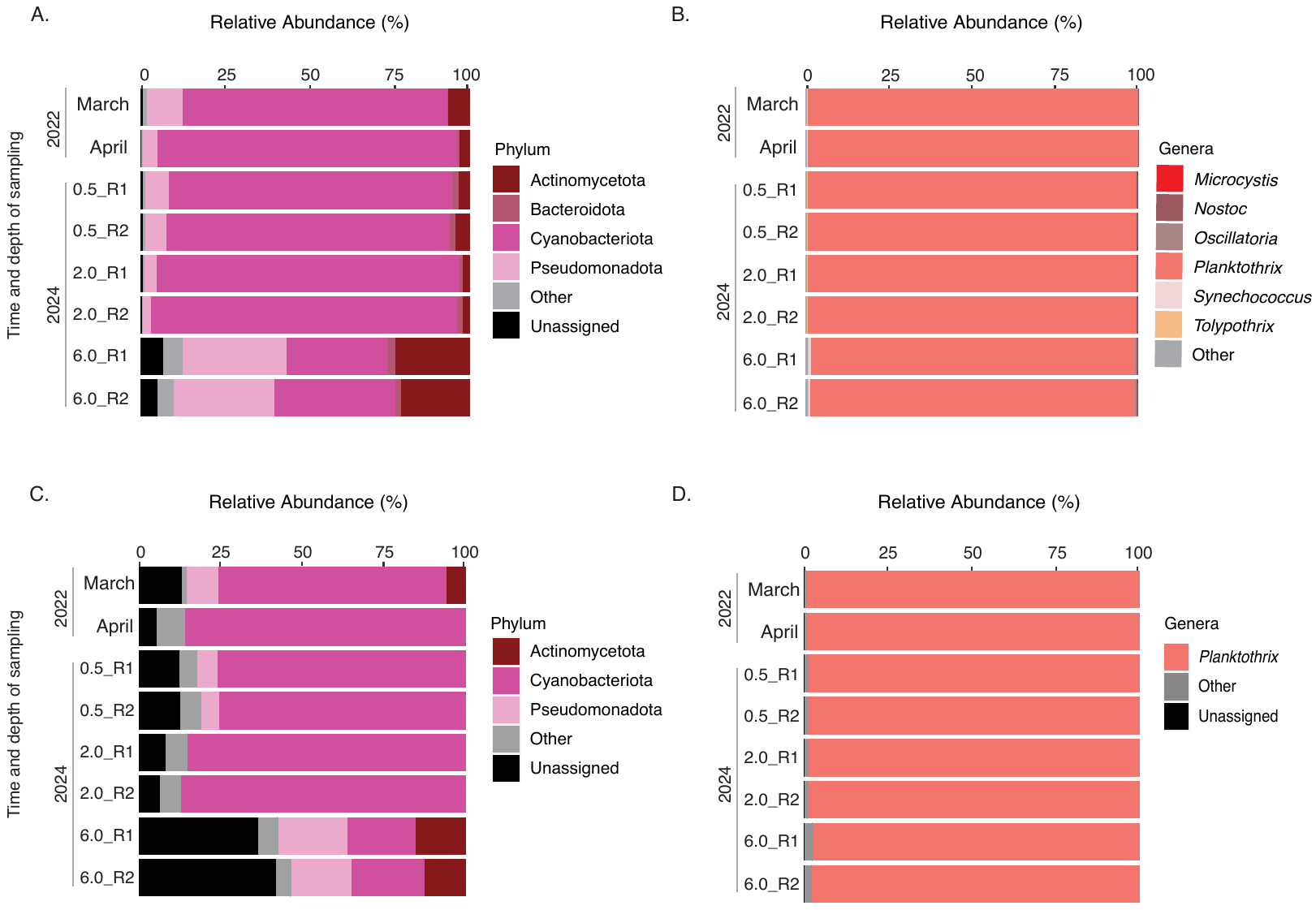

Supplemental Figure 7: Method comparison of two separate approaches to calculating relative abundances of prokaryotic phyla and cyanobacterial genera in Mead’s Quarry. (A-B) Estimates of relative abundance performed *via* manual sorting and curation in Microsoft Excel. Panel A demonstrates relative abundance of major prokaryotic phyla derived from metagenomic libraries, with “Other” defined as phyla contributing <1% of the total reads across all libraries. Panel B demonstrates relative abundance of major cyanobacterial genera derived from metagenomic libraries, with “Other” defined as genera contributing < 0.05% of the total reads across all libraries. (C-D) Estimates of relative abundance performed using phyloseq R package (v.3.20) (McMurdie and Holmes, 2013). Panel A demonstrates relative abundance of major prokaryotic phyla derived from metagenomic libraries, with “Other” defined as phyla contributing < 1% of the total reads across all libraries. Panel B demonstrates relative abundance of major cyanobacterial genera derived from metagenomic libraries, with “Other” defined as genera contributing <1% of the total reads across all libraries.

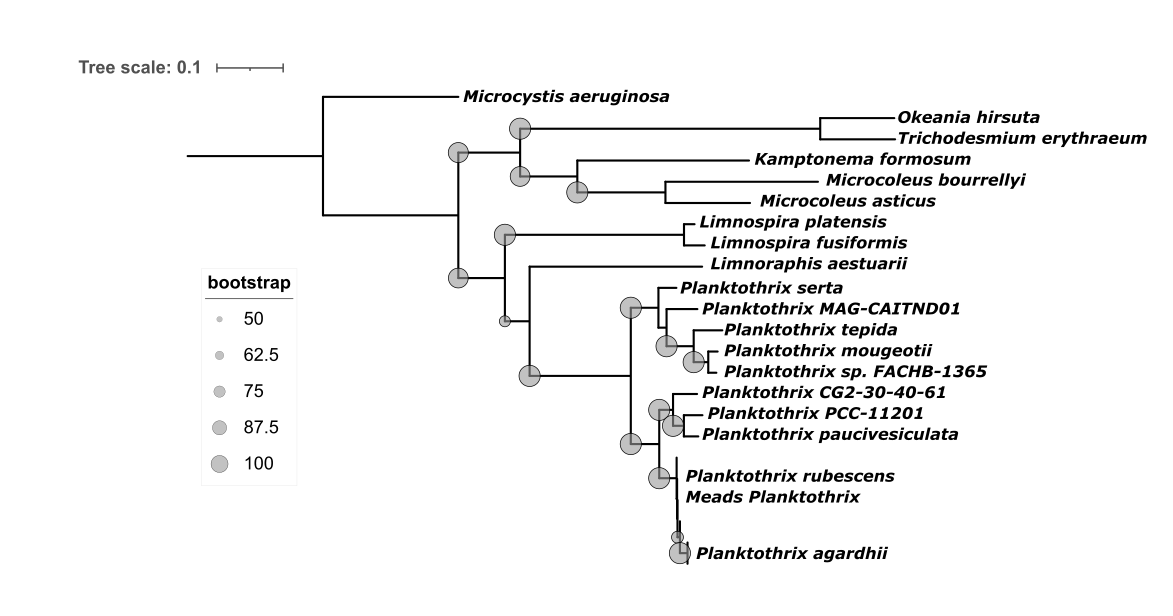

Supplemental Figure 8: Phylogenetic reconstruction of the Microcoleaceae family based on the *rpoB* gene nucleotide sequence. Type representatives of each species as defined by GTDB v. R214 are included. Bootstrap values larger than 50 are annotated at nodes.

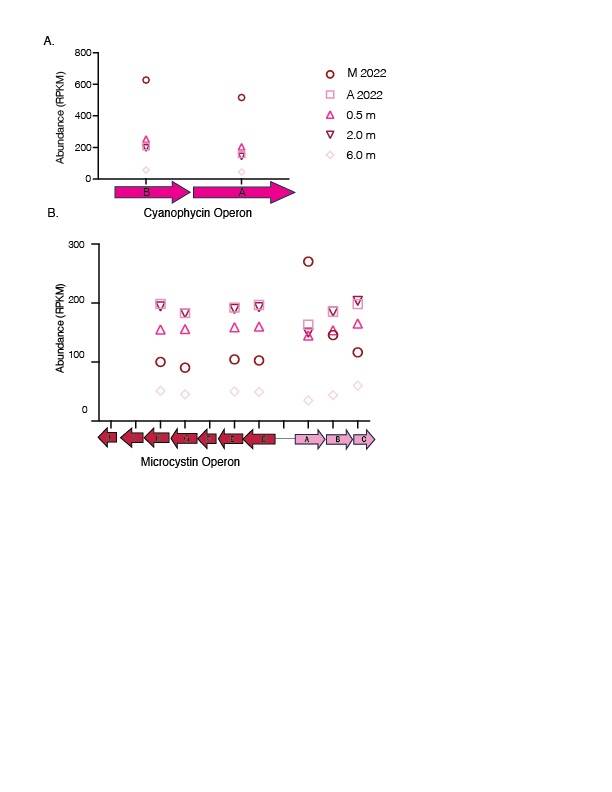

Supplemental Figure 9: (A) Mean relative abundance (RPKM) of genes mapping to the cyanophycin operon in all sample months and sites of interest: March 2022 = dark red circle, April 2022 = bubblegum pink square, 0.5 m depth samples from March 2024 = hot pink triangle, 2.0 m depth samples from March 2024 = dark purple inverted triangle, 6.0 m depth samples from March 2024 = light pink diamond. (B) Mean relative abundance (RPKM) of genes mapping to the microcystin operon in all sample months and site of interest: March 2022 = dark red circle, April 2022 = bubblegum pink square, 0.5 m depth samples from March 2024 = hot pink triangle, 2.0 m depth samples from March 2024 = dark purple inverted triangle, 6.0 m depth samples from March 2024 = light pink diamond

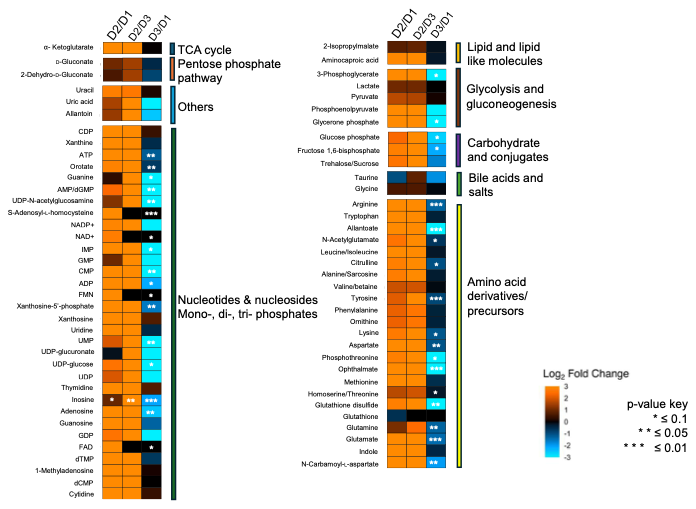

Supplemental Figure 10: Heatmap depicting proportional abundance of detected metabolites organized by compound class between the 0.5 m (D1), 2.0 m (D2) and 6.0 m (D3). The heatmap displays fold differences for comparisons between D2 vs D1; D2 vs D3; and D3 vs D1 (n = 12). Significance of fold change is indicated by asterisks.

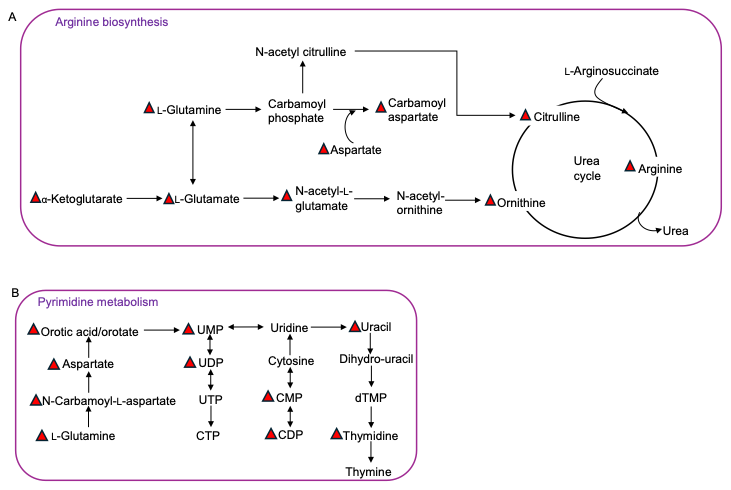

Supplemental Figure 11: Metabolic pathways that were most differentially represented in comparisons between 0.5 m depth and 2.0 m depth. (A) Arginine biosynthesis pathway, with dark red triangles indicating metabolites that were significantly more abundant at 2.0 m compared to 0.5 m depth. (B) Pyrimidine metabolism pathway, with dark red triangles indicating metabolites that were significantly more abundant at 2.0 m compared to 0.5 m depth.
